# Mechanistic Insights of High Temperature-Interfered Meiosis in Autotetraploid *Arabidopsis thaliana*

**DOI:** 10.1101/2021.05.05.442845

**Authors:** Huiqi Fu, Ke Yang, Xiaohong Zhang, Jiayi Zhao, Ibrahim Eid Elesawi, Hong Liu, Jing Xia, Guanghui Yu, Chunli Chen, Chong Wang, Bing Liu

## Abstract

Environmental temperature has a huge impact on multiple meiosis processes in flowering plants. Polyploid plants derived from whole genome duplication are believed to have an enhanced abiotic stress tolerance. In this study, the impact of high temperatures on male meiosis in autotetraploid *Arabidopsis thaliana* was investigated. We found that autotetraploid Columbia (Col-0) plants generate a subpopulation of aberrant meiotic products under normal temperature, which is significantly increased under heat stress. Cytological studies revealed that, as the case in diploid *Arabidopsis thaliana*, assembly of microtubular cytoskeleton network, pairing and segregation of homologous chromosomes, and meiotic recombination in autotetraploid Arabidopsis are compromised under the high temperatures. Immunostaining of ɤH2A.X and recombinase DMC1 suggested that heat stress inhibits formation of DNA double-strand breaks; additionally, it specifically destabilizes ASY1 and ASY4, but not SYN1 on chromosomes. The loading defects of ASY1 and ASY4 overlap in the *syn1* mutant, which supports that the building of lateral element of synaptonemal complex occurs downstream of a SYN1-ASY4-ASY3 stepwise assembly of axis. Remarkably, heat-induced abnormalities of ASY1 and ASY4 co-localize on chromosomes of both diploid and autotetraploid Arabidopsis, suggesting that high temperatures interfere with ASY1-associated SC via an impacted stability of chromosome axis. Furthermore, ZYP1-dependent transverse filament of SC is disrupted by heat stress. Taken together, these findings suggest that polyploidization negatively contributes to instability of chromosome axis and meiotic recombination in *Arabidopsis thaliana* under heat stress.

## Introduction

Meiosis is a specialized type of cell division that, in plants, occurs in pollen mother cells (PMCs) and/or megasporocytes giving rise to gametes with halved ploidy. At early stages of meiosis, meiotic recombination (MR) takes place between homologous chromosomes to drive exchange of genetic information via formation of crossovers (COs). MR results in novel combination of genetic alleles among progenies, which enables natural selection can happen in the population, and safeguards balanced segregation of homologous chromosomes that is vital for production of viable gametes and fertility (Wang and Copenhaver, 2018). MR is initiated by the generation of DNA double-strand breaks (DSBs), which are catalyzed by SPO11, a type-II topoisomerase (topoisomerase VI, subunit A) conserved among eukaryotes (Bergerat et al., 1997; Da Ines et al., 2020; Grelon et al., 2001; Stacey et al., 2006). In Arabidopsis, SPO11-1 and SPO11-2 are required for MR, while SPO11-3 plays a role in endoreduplication (Grelon et al., 2001; Hartung et al., 2007; Stacey et al., 2006; Sugimoto-Shirasu et al., 2002; Yin et al., 2002). Plants with defective DSB formation exhibit impaired homolog synapsis and recombination, and are male sterile due to mis-segregation of chromosomes (Da Ines et al., 2020; De Muyt et al., 2007; Grelon et al., 2001; Stacey et al., 2006; Xue et al., 2018). DSBs are subsequently processed by recombinases RAD51, which repairs DSBs using sister chromatids as a template that leads to non-crossovers (NCOs); or DMC1, which drives MR-specific DSB repair (Da Ines et al., 2013; Klimyuk and Jones, 1997; Kobayashi et al., 2019; Li et al., 2004; Pohl and Nickoloff, 2008; Sanchez-Moran et al., 2007; Singh et al., 2017; Su et al., 2017; Yao et al., 2020). RAD51 and DMC1 do not act independently, with RAD51 functioning as an accessory factor of DMC1 in catalyzing MR (Cloud et al., 2012; Da Ines et al., 2013; Kurzbauer et al., 2012; Lan et al., 2020). There are two types of COs, most of which (∼85%) belong to type-I class catalyzed by ZMM proteins, and are spaced on chromatin by interference (Higgins et al., 2004); while the other COs (type-II) mediated by MUS81 are interference-insensitive (Berchowitz et al., 2007; Hollingsworth and Brill, 2004).

DSB formation and MR rely on a programmed building of chromosome axis. Meiotic-specific cohesion protein AtREC8/SYN1 binds sister chromatids together and aids the chromosomes to form a loop structure (Shahid, 2020; Zickler and Kleckner, 1999). It is proposed that DSBs are formed at the basal region of the loops that are anchored to the ASY1-associated lateral element of synaptonemal complex (SC) (Kim and Choi, 2019; Zickler and Kleckner, 1999). The coiled-coil axis proteins ASY3 and ASY4 play a key role in organizing axis formation, and mediates the connections between the SYN1-mediated chromosome axis and SC via the interplay with ASY1 (Chambon et al., 2018; Ferdous et al., 2012; Osman et al., 2018). Dysfunction of the axis components causes disrupted axis structure and reduced DSB formation, and consequently results in failed homolog synapsis and MR (Bai et al., 1999; Cai et al., 2003; Chambon et al., 2018; Ferdous et al., 2012; Lambing et al., 2020b). SC formation is essentially required for normal homolog synapsis and CO formation. ASY1 contributes to CO formation via the DMC1-mediated MR pathway (Armstrong et al., 2002; Sanchez-Moran et al., 2007). Meanwhile, ASY1 prevents the preferential occurrence of COs at distal regions by antagonizing telomere-led recombination, and by maintaining CO interference along the chromosomes (Lambing et al., 2020a). ZYP1 is a conserved transverse filament protein of SC which is required for homolog synapsis (Barakate et al., 2014; Higgins et al., 2005; Wang et al., 2010). Recent findings revealed that ZYP1-dependent SC formation is indispensable for maintenance of interference, by which ZYP1 restricts the number of type-I COs along the chromosomes; moreover, the bias of CO rate between sexes is wiped when ZYP1 is knocked out (Capilla-Pérez et al., 2021; France et al., 2021). Homologous chromosomes are separated by the bipolar pulling of spindles at the end of meiosis I (MI); and after meiosis II (MII), sister chromatids disjoin with each other, which leads to production of four isolated chromosome sets (Bhatt et al., 2001; Zamariola et al., 2014). In Arabidopsis, like other dicot plants, meiotic cytokinesis takes place thereafter the completion of two rounds of chromosome separation (De Storme and Geelen, 2013).

Male meiosis in plants is sensitive to variations of environmental temperature (Bomblies et al., 2015; De Storme and Geelen, 2014; Liu et al., 2019; Lohani et al., 2019). In both dicots and monocots, low temperatures predominantly affect cytokinesis by disturbing the formation of phragmoplast, which thereby induces meiotic restitution and formation of unreduced gametes (De Storme et al., 2012; Liu et al., 2018; Tang et al., 2011). In contrast, under high temperatures, both chromosome dynamics and cytokinesis are prone to be impacted; especially, the response of MR to heat stress is more complex (De Storme and Geelen, 2020; Draeger and Moore, 2017; Lei et al., 2020; Mai et al., 2019; Ning et al., 2021; Wang et al., 2017). In Arabidopsis, a mild increase of temperature (28°C) positively affects type-I CO rate by enhancing the activity of ZMM proteins without impacting DSB formation (Lloyd et al., 2018; Modliszewski et al., 2018). Under a higher temperature (32°C), however, the rate and distribution of COs are altered (De Storme and Geelen, 2020). Moreover, at extreme high temperatures (36-38°C) over the fertile threshold of Arabidopsis, occurrence of COs is fully suppressed due to inhibited DSBs generation and impaired homolog synapsis; additionally, the microtubular cytoskeleton-based chromosome segregation is disrupted (Lei et al., 2020; Ning et al., 2021). Environmental temperatures therefore may manipulate genomic diversity, and/or influence ploidy consistency of plants over generations by impacting male meiosis during microsporogenesis (Bomblies et al., 2015; Lohani et al., 2019).

Most higher plants, especially for angiosperms, have experienced at least one episode of whole genome duplication (WGD) event, which is considered an important force driving speciation, diversification, and domestication (Del Pozo and Ramirez-Parra, 2015; Dubcovsky and Dvorak, 2007; Leitch and Leitch, 2008; Ren et al., 2018; Soltis et al., 2015). Polyploids are classified into autopolyploids and allopolyploids, which originate from intraspecies WGD events, or arise from multiple evolutionary lineages through the combination of differentiated genomes, respectively (Bretagnolle and Thompson, 1995; Jackson and Chen, 2010; Parisod et al., 2010; Ramsey and Schemske, 1998; Soltis and Soltis, 2009). In autotetraploid plants, four intraspecies-homologues usually undergo randomly separation at anaphase I; this is different as allotetraploids, in which subgenomes tend to segregate independently due to CO formation between the genetically-closer pairs of homologues (Ramsey and Schemske, 2002; Stift et al., 2008). It is believed that the increased sets of homologous chromosomes contribute to genome flexibility and confer the plants with enhanced tolerance to both endogenous genetic mutations, or exogenous environmental stresses (Comai, 2005; Del Pozo and Ramirez-Parra, 2015; Rao et al., 2020; te Beest et al., 2012; Van de Peer et al., 2020; Wu et al., 2020). However, the multiple chromosome sets also challenge genome stability by impacting homolog pairing and balanced chromosome segregation with associated reduced fertility or viability of plants (Comai, 2005; Otto, 2007; Santos et al., 2003; Svačina et al., 2020; Yant et al., 2013). It is proposed that polyploids have evolutionarily developed a moderate strategy that assures genome stability to a large scale by early-stage homoeologous chromosome sorting, chromosome axis-mediated MR modification, and/or by sacrificing an acceptable reduction of CO formation (Bomblies et al., 2016; Grandont et al., 2014; Lloyd and Bomblies, 2016; Morgan et al., 2020; Seear et al., 2020). However, it remains not yet clear how male meiosis in polyploid plants responds to increased environmental temperatures.

In this study, we found that autotetraploid Col-0 plants incubated at 20°C produce a low but consistently-detectable rate of abnormal meiotic products, suggesting that meiotic defects naturally take place in autotetraploid Arabidopsis. We also showed that both the chromosome dynamics and axis formation in the autotetraploid Arabidopsis plants are more sensitive to high temperatures than that in diploid Arabidopsis, suggesting that a duplicated genome does not confer a higher tolerance but instead increases chromosome instability in *Arabidopsis thaliana* under thermal conditions. Remarkably, we provided evidence supporting that ASY1-associated lateral element of SC formation relies on a SYN1-ASY4-ASY3 stepwise assembly of chromosome axis, in which the stability of ASY4- and ASY3-mediated axis bridges the impact of high temperature on SC organization. Overall, these findings provide insights into how high temperatures affect male meiosis in autotetraploid *Arabidopsis thaliana*.

## Results

### Heat stress increases meiotic defects in autotetraploid *Arabidopsis thaliana*

To reveal the impact of heat stress on male meiosis of autotetraploid *Arabidopsis thaliana*, we analyzed tetrad-staged PMCs in heat-stressed autotetraploid Col-0 plants by performing orcein staining (Fig. 1). First, fluorescence in situ hybridization (FISH) using a centromere-specific probe was applied in the autotetraploid Col-0 plants, which showed that somatic cells harbored twenty chromosomes representing that the plants were tetraploid (Supplement Fig. S1). In plants under control temperature, most PMCs (∼95.46%) produced tetrads that subsequently developed into normal-sized microspores with a single nucleus (Fig. 1A; B and E). Interestingly, a low proportion of polyads and/or unbalanced-tetrads was consistently observed, which resulted in formation of microspores with varied sizes (Fig. 1A; C, polyad, 3.67%; D, unbalanced-tetrad, 0.86%; F, microspores). These phenotypes suggested that a minor frequency of meiotic defects naturally take place in autotetraploid *Arabidopsis thaliana*. In plants stressed by 37°C, a significantly increased frequency (∼92.49%) of abnormal meiotic products was observed, in which unbalanced-triads, polyads and unbalanced-tetrads occupied the highest proportions (Fig. 1A; M-O, unbalanced-triads, 29.02%; R and S, polyads, 26.42%; P and Q, unbalanced-tetrads, 26.17%). These figures, together with the observed unbalanced-dyads that contained differently-numbered and/or -sized nuclei (Fig. 1A, G, I and J, 7.25%), indicated that chromosome segregation in MI and/or II was interfered. Meanwhile, the occurrence of balanced-dyad and balanced-triad represented an induction of meiotic restitution (Fig. 1A; H, balanced-dyad, 2.85%; K, balanced-triad, 0.78%). The defective tetrad stage PMCs leaded to generation of aneuploid microspores at unicellular stage (Fig. 1T and U). Similar cellular alterations were observed in autotetraploid Col-0 plants stressed by 32°C (Fig. 1A; V-Z). Moreover, we found that under high temperatures, the flower buds with the same size as that in control occurring meiosis and/or cytokinesis contained unicellular stage microspores, which hinted that the high temperatures accelerated meiosis progressing (Supplement Fig. S2A-C).

**Figure 1.**
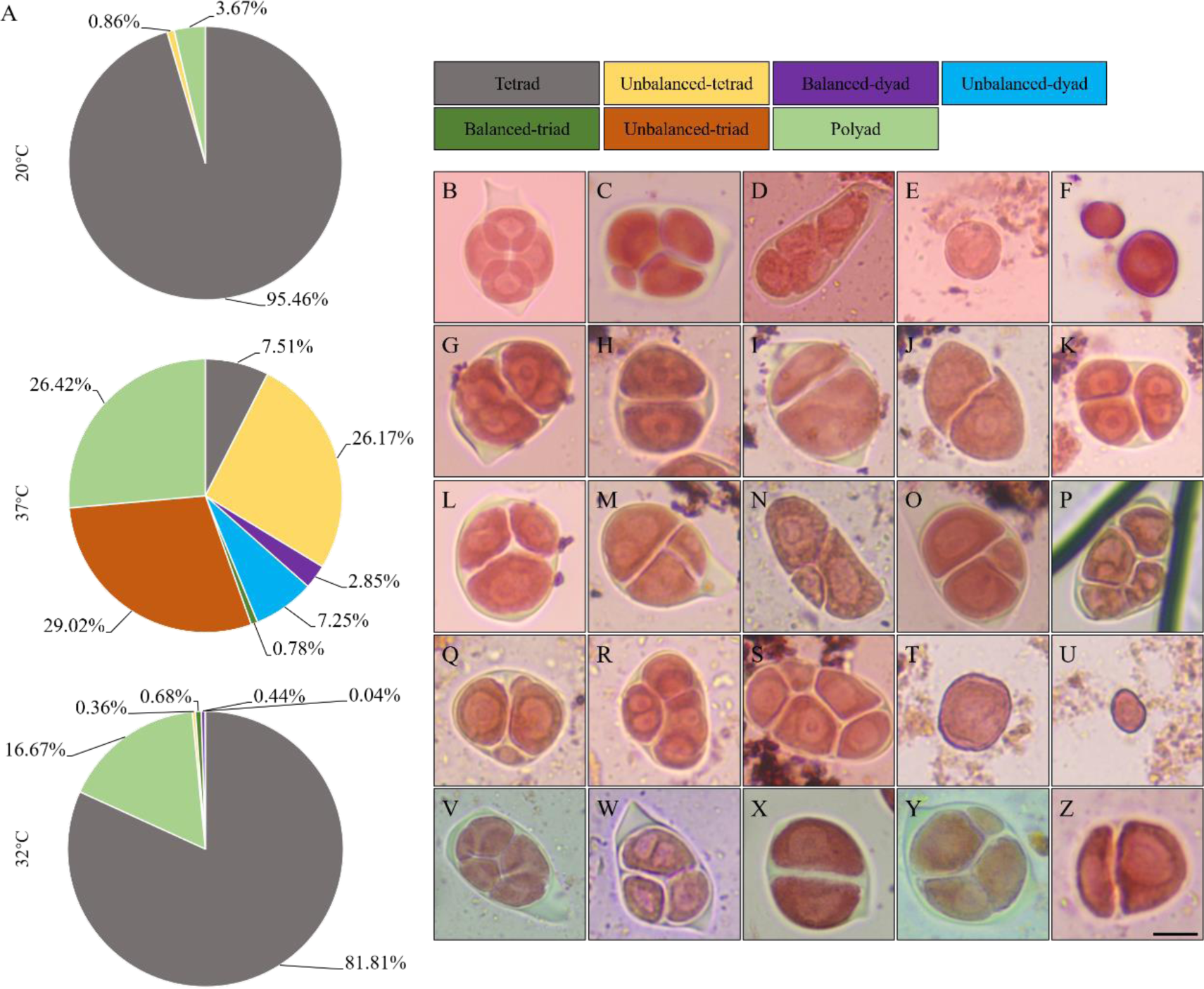
Tetrad analysis in heat-stressed autotetraploid Col-0 plants. A, Graph showing the frequency of tetrad-staged meiotic products in autotetraploid Col-0 plants incubated under 20°C, 32°C and 37°C, respectively. The numbers indicate the frequency of different types of tetrad-staged meiocytes. B-F, Tetrad-staged PMCs (B-D) and unicellular-staged microspores (E and F) of autotetraploid Col-0 plants grown under 20°C. G-U, Tetrad-staged PMCs (G-S) and unicellular-staged microspores (T and U) of autotetraploid Col-0 plants stressed by 37°C. V-Z, Tetrad-staged PMCs from autotetraploid Col-0 plants stressed by 32°C. Scale bar = 10 μm.

We next stained meiotic cell walls using aniline blue to examine the impact of high temperatures on meiotic cytokinesis in autotetraploid Col-0 plants. In line with the analysis by orcein staining, most tetrads in control plants showed a regular ‘cross’-like cell wall formation (Fig. 2A). Observation of polyad supported that meiotic defects naturally occurred in the autotetraploid Arabidopsis (Fig. 2B). After incubation under 37°C, the autotetraploid Col-0 plants showed defective meiotic cell wall formation including unbalanced-triads (Fig. 2C; Supplement Fig. S3F-I), polyad (Fig. 2D; Supplement Fig. S3K and L), unbalanced-tetrads (Fig. 2E; Supplement Fig. S3E and J), unbalanced-dyad (Fig. 2F; Supplement Fig. S3B-D), balanced-dyad (Fig. 2G; Supplement Fig. S3A) and balanced-triad (Fig. 2H; Supplement Fig. S3E). These data demonstrated that heat stress interferes with one or more meiosis processes; e.g. chromosome segregation and cytokinesis in autotetraploid *Arabidopsis thaliana*.

**Figure 2.**
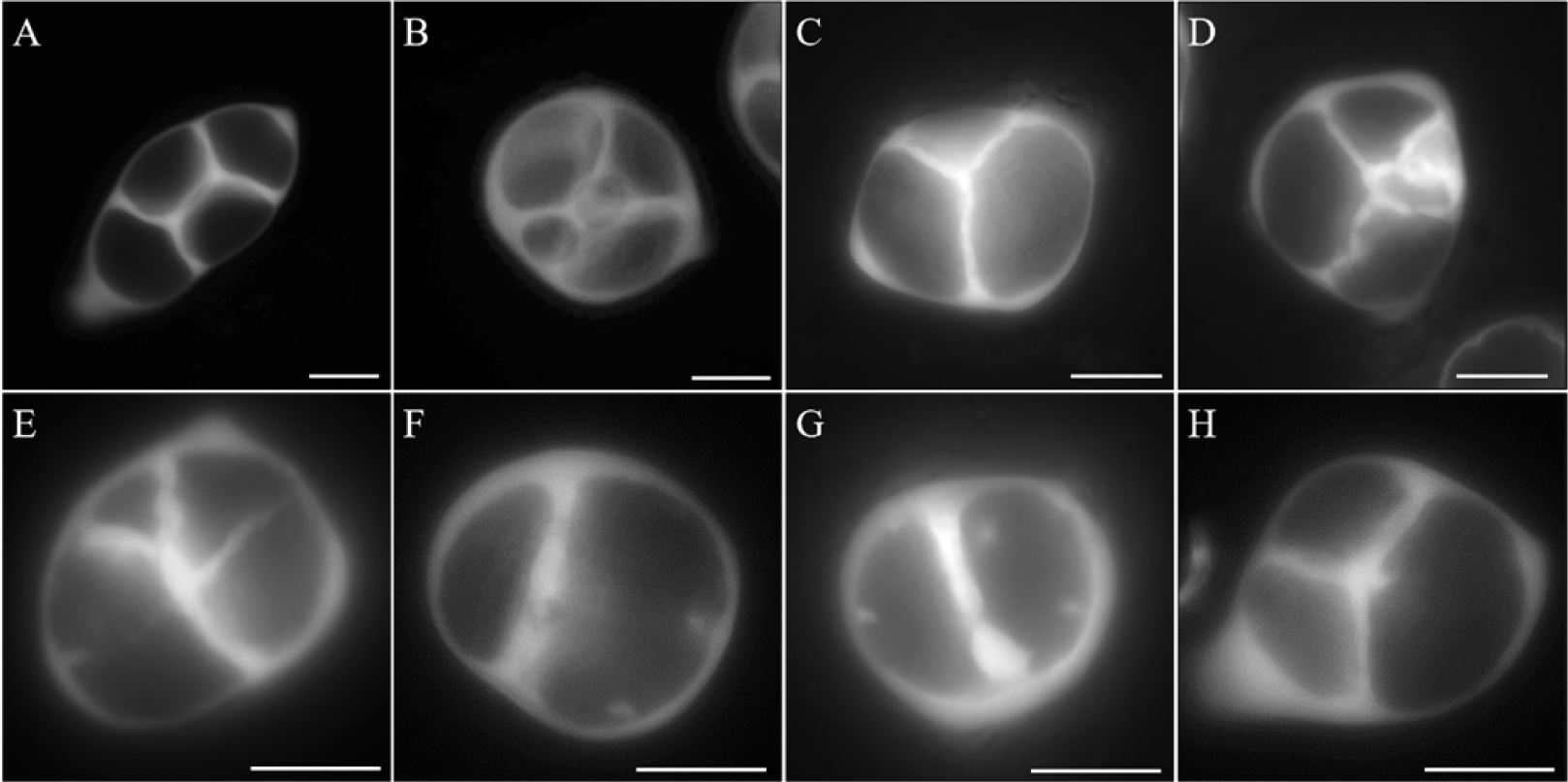
Meiotic cell wall formation in heat-stressed autotetraploid Col-0 plants. A-H, Aniline blue-stained callosic cell walls in autotetraploid Col-0 plants incubated under control (A and B) and high temperature (C-H). Scale bars = 10 μm.

### Microtubular cytoskeleton in autotetraploid Arabidopsis is abnormally assembled under heat stress

To address the cellular mechanism underlining heat-induced aberrant tetrad formation and defective cytokinesis in autotetraploid Arabidopsis, we examined microtubular cytoskeleton by performing immunostaining of α-tubulin (Fig. 3). In autotetraploid Col-0 plants incubated at 20°C, formation of spindle was initiated at early metaphase I (Fig. 3A), and thereafter at middle metaphase I, homologous chromosomes aligned at the cell plate when microtubule arrays got attachment with the centromeres labeled by an anti-CENH3 antibody (Fig. 3B). The monopolar pulling force from the spindle separated the homologous chromosomes at the end of MI, and two spindles were formed at metaphase II to separate the sister chromatids (Fig. 3C). At telophase II, mini-phragmoplast structures composed of radial microtubule arrays (RMAs) were constructed between the four isolated nuclei (Fig. 3D). Notably, triad- and poly-like configurations together with the occurrence of mini-nucleus (Fig. 3E and F) supported existence of meiotic defects in control autotetraploid Arabidopsis plants. After heat treatment, we found that the metaphase I microtubule arrays did not display a typical spindle configuration, which bond with randomly-distributed univalent chromosomes (Fig. 3G and H). The univalents and the impaired spindle leaded to unbalanced segregation of homologous chromosomes at interkinesis, which, meanwhile, showed irregular and sparse phragmoplast formation (Fig. 3I). Disrupted spindles were also observed at metaphase II (Fig. 3J). These alterations resulted in generation of unbalanced-tetrad and polyad with omitted and/or abnormally-shaped RMAs between the adjacent and/or differently-sized nuclei (Fig. 3K and L).

**Figure 3.**
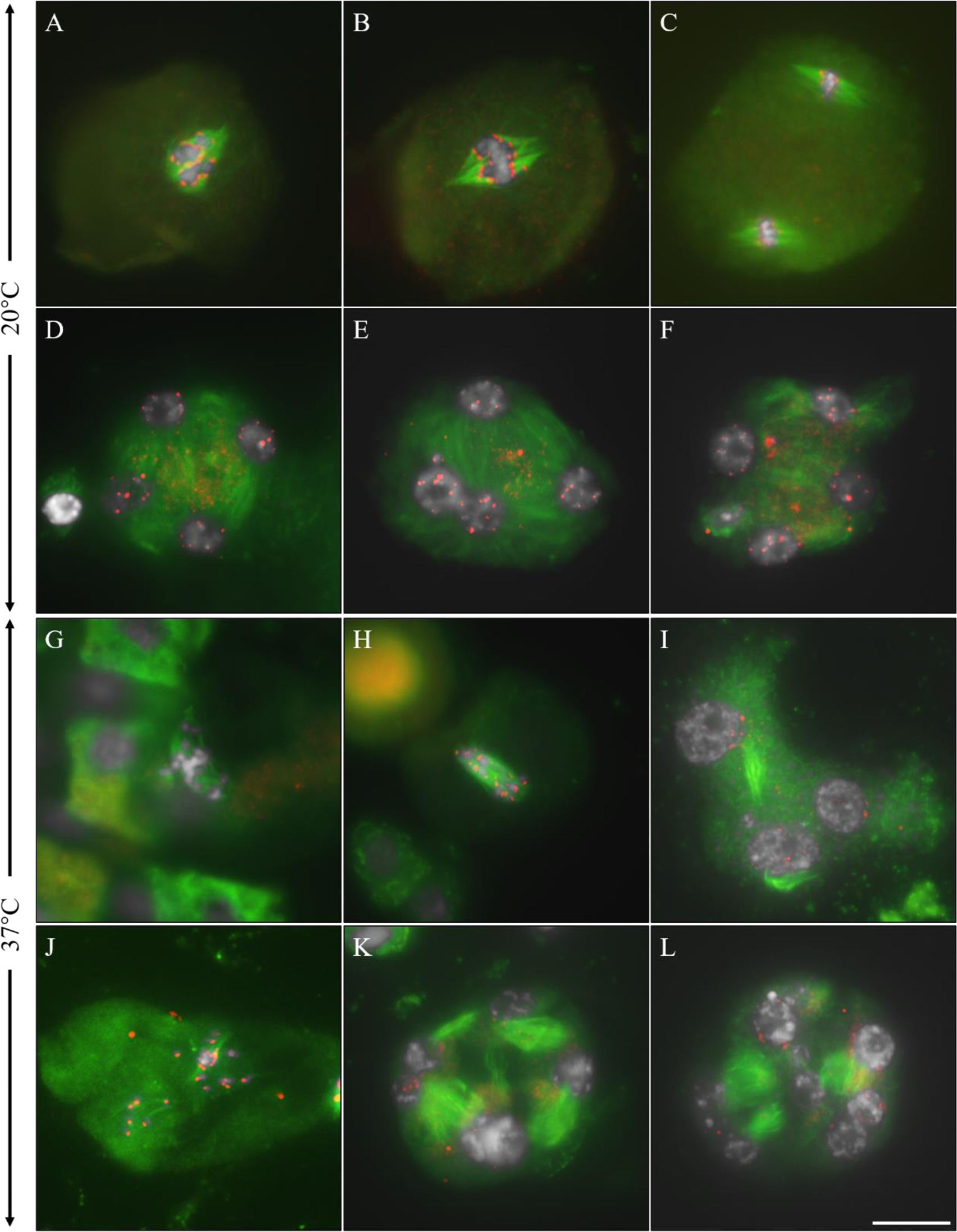
Microtubular cytoskeleton in meiocytes of autotetraploid Col-0 plants. A-F, Metaphase I- (A and B), metaphase II- (C) and telophase II-staged (D-F) PMCs in autotetraploid Col-0 plants under control temperature. G-L, Metaphase I- (G and H), interkinesis-(I), metaphase II- (J) and telophase II-staged (K and L) PMCs in autotetraploid Col-0 plants under high temperature. White, DAPI; green, α-tubulin; red, CENH3. Scale bar = 10 μm.

### Heat stress interferes with chromosome behaviors in autotetraploid *Arabidopsis thaliana*

Chromosome behaviors in meiocytes of autotetraploid Col-0 plants were analyzed using 4′,6-diamidino-2-phenylindole (DAPI) staining. In control plants, homologous chromosomes synapsed at pachytene (Fig. 4A). However, unpaired chromosomes were occasionally observed indicating a pairing defect (Fig. 4B). Bridge- and thick rope-like structures implied irregular chromosome interactions (Fig. 4C and D, yellow and green arrows). At diakinesis and metaphase I, most meiocytes showed existence of five tetravalents (Fig. 5E-G and I; 60.71%, n = 34). About 33.93% meiocytes showed a combination of bivalents and tetravalents including PMCs containing four (21.43%, n = 12), three (8.93%, n = 5) and two tetravalents (3.57%, n = 2) (Fig. 4H, J and K, red arrow). Additionally, we observed meiocytes that harbored ten bivalents (Fig. 3L; 5.36%, n = 3). These suggested that synapsis took place between four homologues or a couple of them, or occurred only between two homologous chromosomes. Meanwhile, irregular connections hinted associations between nonhomologous chromosomes (Fig. 4F, G and J, yellow arrows). After MI, most meiocytes underwent balanced segregation of the four chromosome sets (Supplement Fig. S4A and D; 75%, n = 48), which resulted in generation of tetrads with four isolated nuclei each harboring a halved chromosome number (Supplement Fig. S4G; 88.24%, n = 105). Unbalanced separation of homologues with lagged chromosomes (Supplement Fig. S4B and C, E and F; 25%, n = 16) consequently leaded to defective tetrads (Supplement Fig. S4H-J; 11.76%, n = 14).

**Figure 4.**
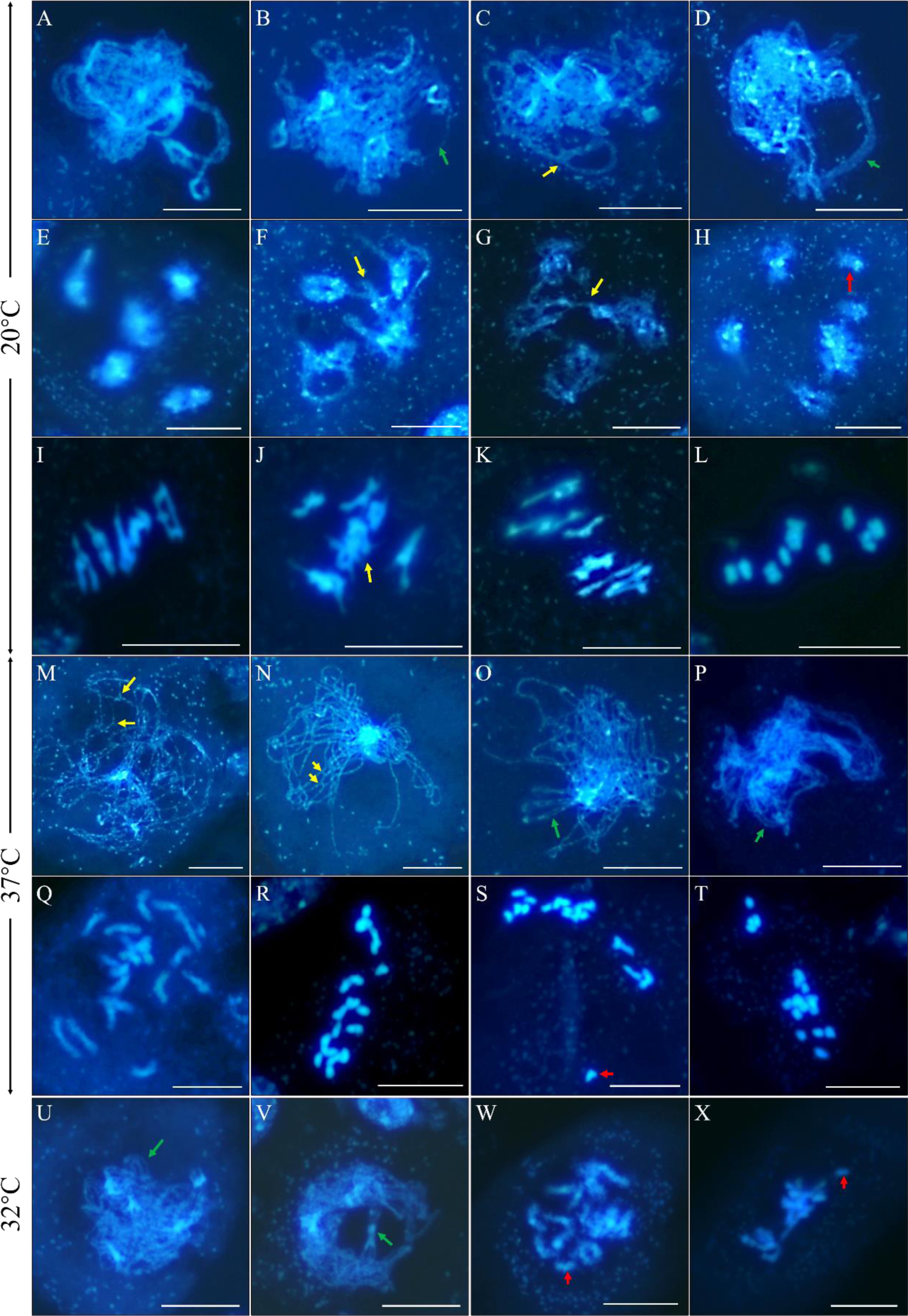
Chromosome behaviors in meiocytes of autotetraploid Col-0 plants. A-L, Chromosome spreading of Pachytene- (A-D), diakinesis- (E-H) and metaphase I-staged (I-L) meiocytes in autotetraploid Col-0 plants grown under control temperature. M-T, Chromosome spreading of zygotene- (M and N), pachytene- (O and P), diakinesis- (Q) and metaphase I-staged (R-T) meiocytes in autotetraploid Col-0 plants stressed by 37°C. U-X, Chromosome spreading of pachytene- (U and V), diakinesis- (W) and metaphase I-staged (X) meiocytes in autotetraploid Col-0 plants stressed by 32°C. Green arrows indicate incomplete and/or abnormal pairing of chromosomes; yellow arrows indicate abnormal chromosome interactions; and red arrows indicate bivalents, univalents and/or lagged chromosomes. Scale bars = 10 μm.

**Figure 5.**
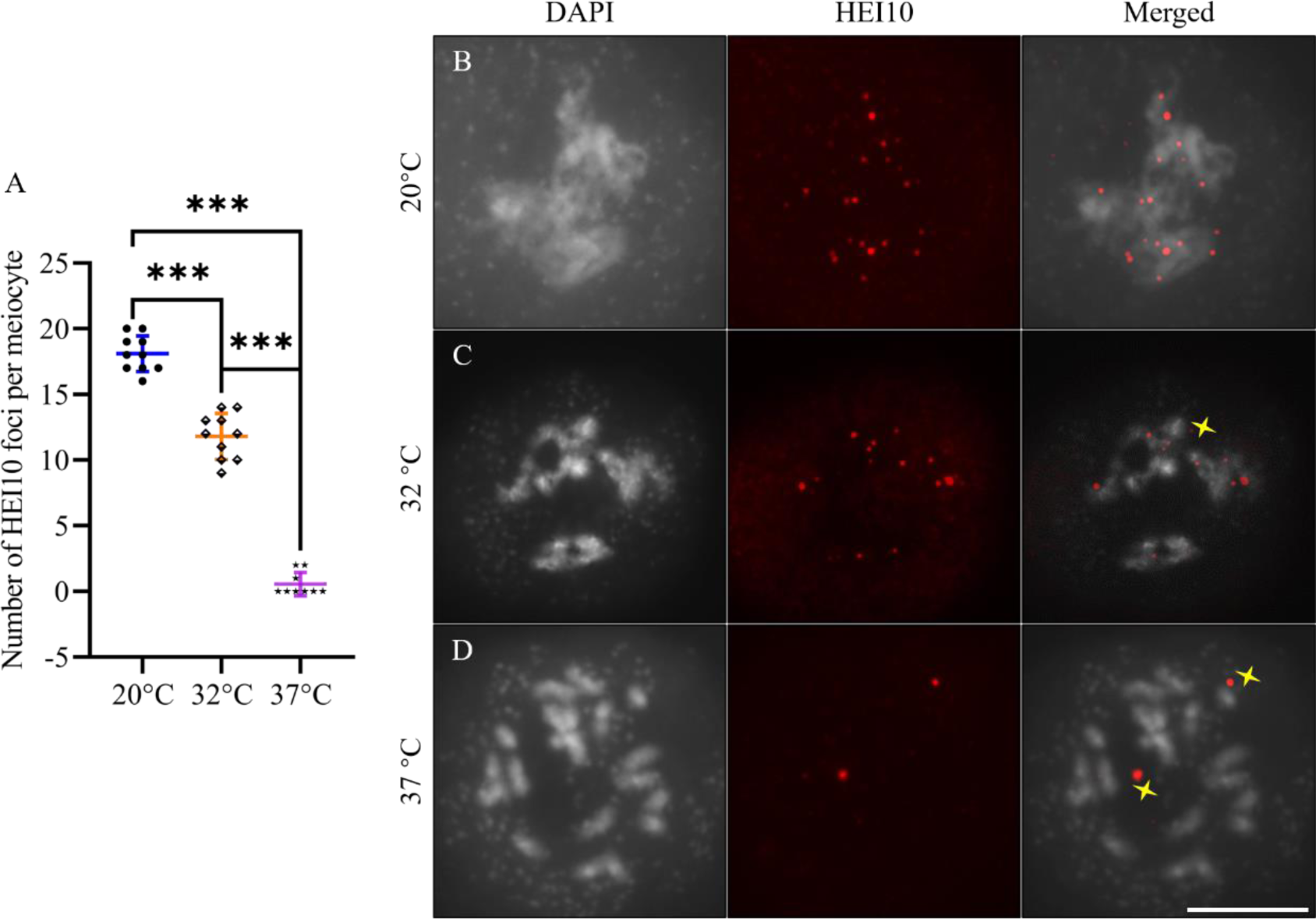
Localization of HEI10 on diakinesis chromosomes in autotetraploid Col-0 plants. A, Graph showing the number of HEI10 foci per diakinesis-staged meiocyte. One-way ANOVA test was performed, and the significance level was set as *P* < 0.05. *** indicates *P* < 0.0001. B-D, Immunolocalization of HEI10 on diakinesis chromosomes of autotetraploid Col-0 plants incubated under 20°C (B), 32°C (C) and 37°C (D), respectively. The yellow stars indicate non-specific foci to HEI10. Scale bar = 10 μm.

In autotetraploid Col-0 plants treated by 37°C, we observed bridge-like chromosome structures at zygotene (Fig. 4M and N, yellow arrows); meanwhile, incomplete and/or impaired pairing and synapsis were found on the pachytene chromosomes (Fig. 4O and P, green arrows). These figures suggested that the high temperature induced abnormal chromosome interactions and interfered with homolog synapsis. At diakinesis and metaphase I, ∼93.68% meiocytes showed twenty individual chromatids (Fig. 4Q-T, red arrow; n = 89), which represented an omitted and/or suppressed CO formation that consequently resulted in unbalanced chromosome segregation at anaphase I (98.08%, n = 51) and tetrad stage (97.67%, n = 126) (Supplement Fig. S4K-T). Similarly, in autotetraploid Col-0 plants stressed under 32°C, failed and/or irregular chromosome pairing and synapsis were induced (Fig. 4U and V, green arrows); in the meantime, a combination of univalents, bivalents and multivalents suggested that CO formation was compromised (Fig. 4W and X, red arrows) which leaded to unbalanced chromosome segregation at MII (Supplement Fig. S4U-Y). Taken together, these findings revealed that heat stress interferes with chromosome pairing and/or synapsis, and segregation in autotetraploid Arabidopsis.

### High temperatures reduce abundance of HEI10 on diakinesis chromosomes

Increased temperature influences MR rate by modulating type-I CO formation in Arabidopsis (Modliszewski et al., 2018). Univalents in heat-stressed autotetraploid Col-0 plants suggested a reduced CO rate under the high temperatures. To this end, we quantified the number of HEI10 that acts in type-I CO formation pathway in diakinesis-staged meiocytes of autotetraploid Col-0 plants. The plants grown under control temperature showed ∼18.10 HEI10 foci per meiocyte (Fig. 5A and B); by contrast, the abundance of HEI10 reduced to ∼11.80 and ∼0.56 per meiocyte in the plants stressed by 32°C and 37°C, respectively (Fig. 5A, C and D), which suggested that the high temperatures significantly inhibited occurrence of type-I class CO in the autotetraploid Col-0 plants.

### DSB formation in autotetraploid Arabidopsis is suppressed under high temperatures

Generation of DSB is crucial for CO formation (De Muyt et al., 2007; Hartung et al., 2007; Kurzbauer et al., 2012). To test whether heat-induced reduction of CO was owing to a compromised DSB formation, like in diploid Arabidopsis (Ning et al., 2021), we quantified the number of ɤH2A.X, which specifically marks DSB sites, on zygotene chromosomes. Under control temperature, an average of ∼146.5 ɤH2A.X foci per meiocyte was detected (Fig. 6A and D). In plants stressed by 32°C and 37°C, however, the abundance of ɤH2A.X was reduced to ∼79.5 and ∼84.8 per meiocyte, respectively (Fig. 6B-D), which indicated a significantly lowered DSB formation. In support of this, the number of DMC1 that specifically catalyzes DSB repair for MR was reduced to ∼39.4 and ∼49.7 per meiocyte under 32°C and 37°C, respectively, which were much lowered compared with control (∼142) (Fig. 6E-H). These data provided evidence that high temperatures impose a negative impact on DSB formation in autotetraploid Arabidopsis.

**Figure 6.**
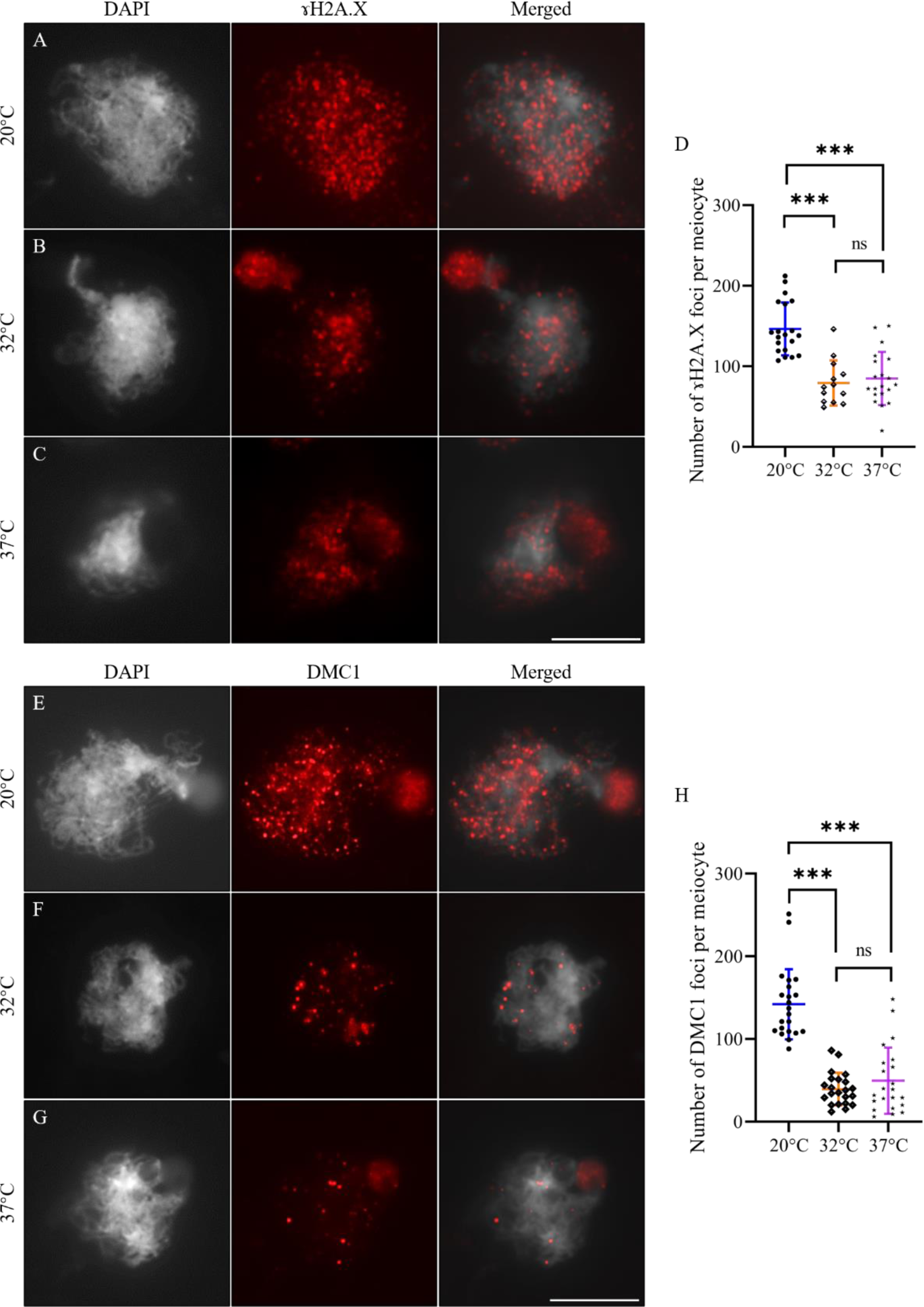
Heat stress reduces abundance of ɤH2A.X and DMC1 on zygotene chromosomes. A-C, Immunolocalization of ɤH2A.X on zygotene chromosomes of autotetraploid Col-0 plants under 20°C (A), 32°C (B) and 37°C (C), respectively. D, Graph showing the number of ɤH2A.X foci per meiocyte in autotetraploid Col-0 plants under 20°C, 32°C and 37°C. E-G, Immunolocalization of DMC1 on zygotene chromosomes of autotetraploid Col-0 plants under 20°C (E), 32°C (F) and 37°C (G), respectively. H, Graph showing the number of DMC1 foci per meiocyte in autotetraploid Col-0 plants under 20°C, 32°C and 37°C. One-way ANOVA test was performed, and the significance level was set as *P* < 0.05. *** indicates *P* < 0.001; ns indicates *P* > 0.05. Scale bars = 10 μm.

### Heat stress destabilizes ASY1 and ASY4 accumulation on chromosomes

To reveal the impact of high temperature on chromosome axis formation in autotetraploid Arabidopsis, we analyzed loading of the main axis-associated components; i.e. SYN1, ASY1 and ASY4 proteins in heat-stressed autotetraploid Col-0 plants. Under control temperature, linear SYN1 and ASY1 signals overlapped and were fully associated with the entire zygotene chromosomes (Fig. 7A). At late pachytene, when homologous chromosomes synapsed, ASY1 were unloaded at some chromosome regions (Fig. 7B). After heat treatment, although that some zygotene chromosomes showed uninfluenced accumulation of ASY1 (Fig. 7C and D; Supplement Fig. S5A, 19.55%; Supplement Fig. S6A), most zygotene and pachytene chromosomes displayed dotted configuration of ASY1 (Fig. 7E and F, yellow arrows; Supplement Fig. S5A, 80.45%; Supplement Fig. S6B-F), suggesting an impacted stability of ASY1-associated axis. By contrast, no obvious alteration in SYN1 loading was observed in heat-stressed meiocytes (Fig. 7C-F, I-K).

**Figure 7.**
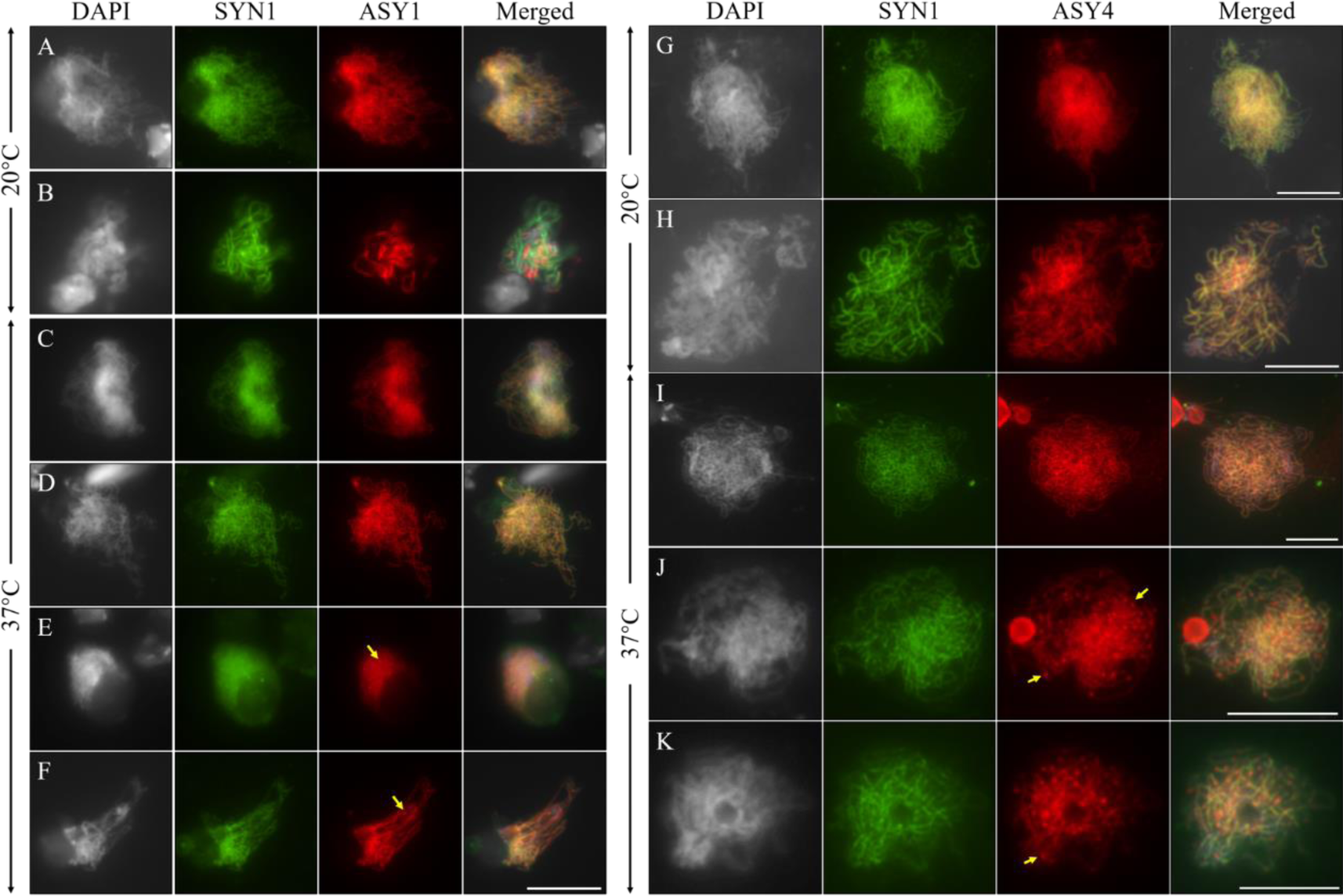
Immunostaining of chromosome axis components in meiocytes of autotetraploid Col-0 plants. A and B, Co-immunostaining of SYN1 and ASY1 on zygotene (A) and pachytene (B) chromosomes in control plants. C-F, Co-immunostaining of SYN1 and ASY1 on zygotene (C-E) and pachytene (F) chromosomes in heat-stressed plants. G and H, Co-immunostaining of SYN1 and ASY4 on zygotene (G) and pachytene (H) chromosomes in control plants. I-K, Co-immunostaining of SYN1 and ASY4 on zygotene (I and J) and pachytene (K) chromosomes in heat-stressed plants. Yellow arrows indicate dotted ASY1 and ASY4 foci. Scale bars = 10 μm.

On the other hand, early zygotene chromosomes in autotetraploid Col-0 displayed thin linear ASY4 loading (Supplement Fig. S7A), which was extended to the whole chromosomes from middle zygotene to middle pachytene (Fig. 7G and H; Supplement Fig. S7B and C). As the progressing of MR, ASY4 signals showed disassociation with some chromosome regions from late pachytene (Supplement Fig. S7D), and were further unloaded at diplotene with remaining dotted ASY4 foci on diakinesis-staged chromosomes (Supplement Fig. S7E-G). In heat-stressed plants, a minor proportion of zygotene and pachytene chromosomes displayed normal ASY4 accumulation (Fig. 7I; Supplement Fig. S5B, 19.85%; Supplement Fig. S7H and I); on the contrary, a majority of zygotene- and pachytene-staged meiocytes showed dotted ASY4 foci (Fig. 7J and K, yellow arrows; Supplement Fig. S5B, 80.15%; Supplement Fig. S7J and K). Punctate ASY4 signals also occurred on the univalent chromosomes (Supplement Fig. S7L). These findings suggested that heat stress specifically destabilizes the ASY1- and ASY4- but not SYN1-mediated chromosome axis.

### Overlapped abnormalities of ASY1 and ASY4 in the *syn1* mutant

It was proposed that assembly of ASY1-associated lateral element of SC relies on a step-wise formation of SYN1-ASY3-mediated chromosome axis, which is bridged by ASY4 (Chambon et al., 2018; Ferdous et al., 2012; Lambing et al., 2020b). To consolidate the role of ASY4 in mediating ASY1 assembly, we performed co-immunostaining of ASY1 and ASY4 in the *syn1* mutant. At middle zygotene, when homologous chromosomes partially paired and synapsed, ASY1 and ASY4 were fully assembled along the whole chromosomes in the diploid Col-0 plants (Fig. 8A). By contrast, incomplete and/or fragmented ASY1 and ASY4 configuration were observed in zygotene and pachytene meiocytes of the *syn1* mutant (Fig. 8B and C), which, in addition, overlapped (Fig. 8B and C). This supported the notion that ASY1-associated SC assembly relies on ASY4-mediated axis formation, which in turn depends on functional SYN1.

**Figure 8.**
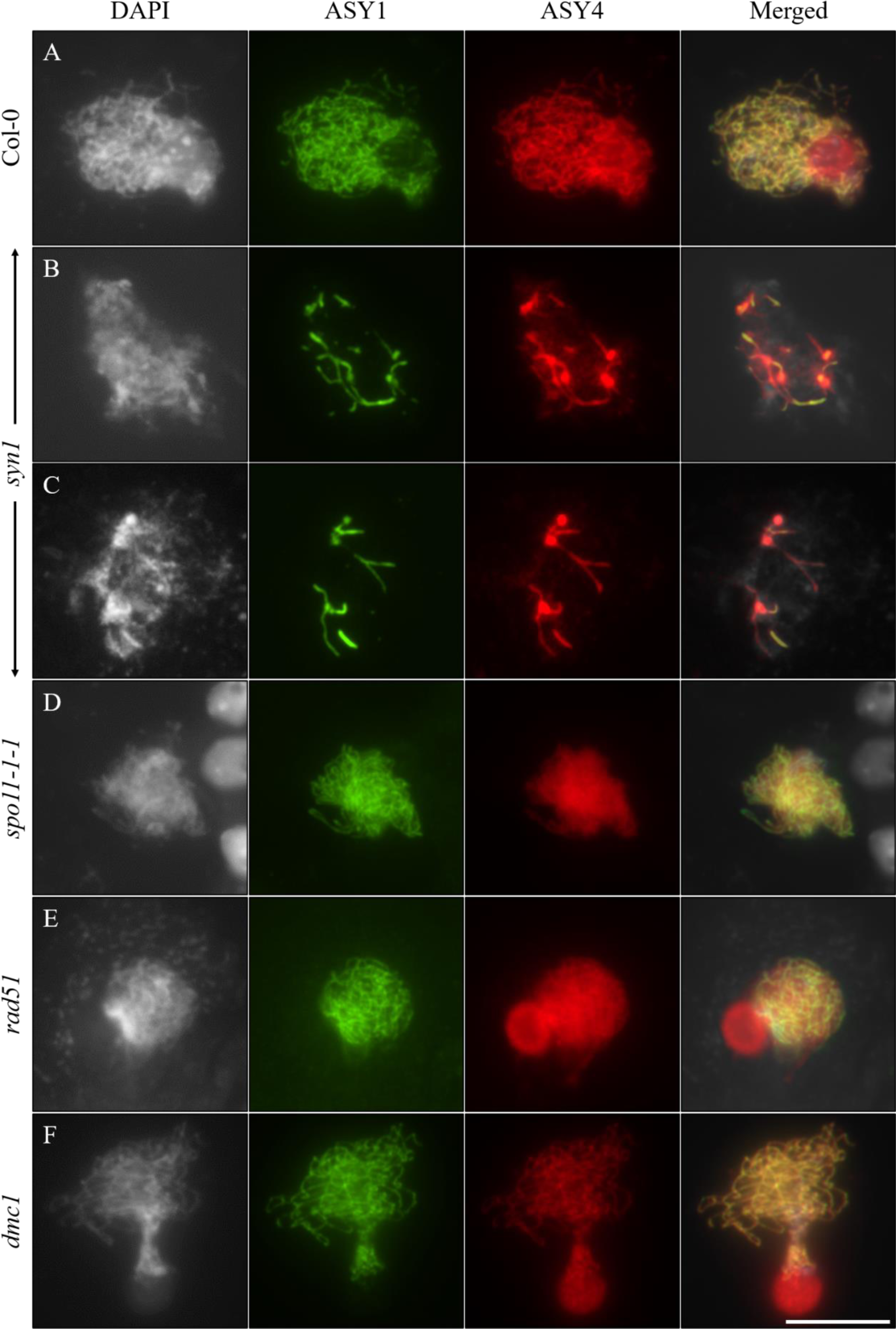
Immunostaining of ASY1 and ASY4 in meiocytes of diploid *Arabidopsis thaliana* plants. A-F, Immunolocalization of ASY1 and ASY4 on prophase I chromosomes of diploid Col-0 plants (A), and the *syn1* (B and C), *spo11-1-1* (D), *rad51* (E) and *dmc1* (F) mutants. Scale bar = 10 μm.

Moreover, DSB formation is considered a downstream event of axis formation and occurs independently of SC assembly (Lambing et al., 2020b; Sanchez-Moran et al., 2007). To consolidate this model, we checked localization of ASY1 and ASY4 in the meiocytes of *spo11-1-1*, *rad51* and *dmc1* mutants. We did not detect any alteration in loading of ASY1 and/or ASY4 on chromosomes of these mutants (Fig. 8D, *spo11-1-1*; E, *rad51* and F, *dmc1*).

### Co-localization of abnormal ASY1 and ASY4 signals on chromosomes of heat-stressed diploid and autotetraploid Arabidopsis

Considering the similarities of defective ASY1 and ASY4 accumulation under heat stress, and the upstream action of axis formation on SC assembly (Fig. 7E and F, J and K; Fig. 8B and C) (Ferdous et al., 2012; Lambing et al., 2020b; Ning et al., 2021), we hypothesized that heat stress destabilizes ASY1-associated SC via impacted ASY4-mediated chromosome axis. To this end, we conducted a combined staining of ASY1 and ASY4 in both the heat-stressed diploid and autotetraploid Col-0 plants. Under control temperature, ASY1 and ASY4 co-localized on the entire chromosomes of diploid and autotetraploid Col-0 at early and middle zygotene (Fig. 9A and D; Supplement Fig. S8A and B). ASY1 subsequently started to be disassociated with the chromosomes from early to late pachytene, when ASY4 displayed relatively stable linear configuration (Supplement Fig. S8C-E). Dotted ASY1 foci occurred at diplotene, which became sparser at diakinesis representing a completed functioning of ASY1 in mediating homolog synapsis (Supplement Fig. S8F and G). ASY4, however, displayed a slower unloading off the chromosomes, which supported its role in aiding the assembly of ASY1 (Supplement Fig. S8F and G). Interestingly, in heat-stressed diploid and autotetraploid Col-0 plants, incomplete and/or dotted ASY1 and ASY4 signals co-localized on the chromosomes (Fig. 9B and C, E and F, yellow arrows). These observations favored the hypothesis that heat stress destabilizes ASY1-associated SC via a compromised stability of ASY4-mediated axis.

**Figure 9.**
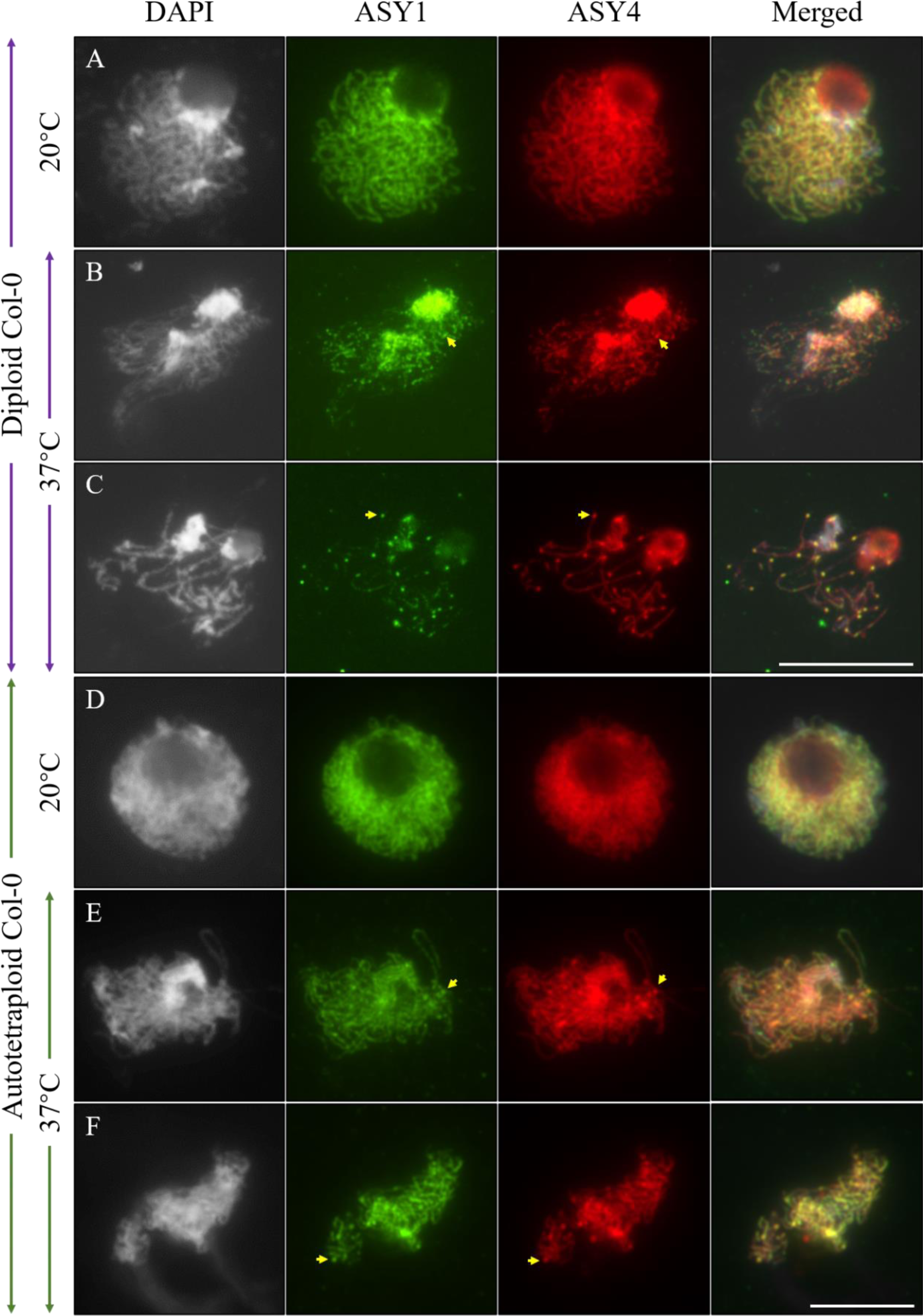
Co-localization of ASY1 and ASY4 in heat-stressed diploid and autotetraploid Col-0 plants. A and D, Zygotene-staged meiocytes in diploid (A) and autotetraploid (D) Col-0 plants grown under control temperature. B and C, Zygotene- (B) and pachytene-staged (C) meiocytes in heat-stressed diploid Col-0 plants. E and F, Zygotene- (E) and pachytene-staged (F) meiocytes in heat-stressed autotetraploid Col-0 plants. Yellow arrows indicate co-localization of dotted ASY1 and ASY4 foci. Scale bars = 10 μm.

### Impaired ZYP1-dependent transverse filament under heat stress

Formation of ZYP1-dependent transverse filament (TF) of SC is required for homolog synapsis (Barakate et al., 2014; Capilla-Pérez et al., 2021; France et al., 2021; Higgins et al., 2005; Wang et al., 2010). We analyzed ZYP1 assembly to examine the impact of heat stress on the central element of SC in autotetraploid Col-0 plants. In control, ZYP1 proteins were linearly and partially loaded at the central regions of paired homologous chromosomes from zygotene (Fig. 10A), which were fully assembled at middle pachytene representing a matured SC formation (Fig. 10B). As the disintegration of SC from late pachytene, ZYP1 proteins were gradually disassociated with chromosomes (Fig. 10C). After heat treatment, dotted and/or fragmented installation of ZYP1 on chromosomes were observed from early zygotene to late pachytene (Fig. 10D-H, yellow arrow), indicating that assembly of transverse filament of SC was impaired. Meanwhile, aggregated and/or enlarged ZYP1 foci implied pairing of multiple chromosomes (Fig. 10D, G and H, blue arrows) (Morgan et al., 2017). Overall, these figures suggested that heat stress disrupts the building of ZYP1-dependent transverse filament of SC in autotetraploid Arabidopsis.

**Figure 10.**
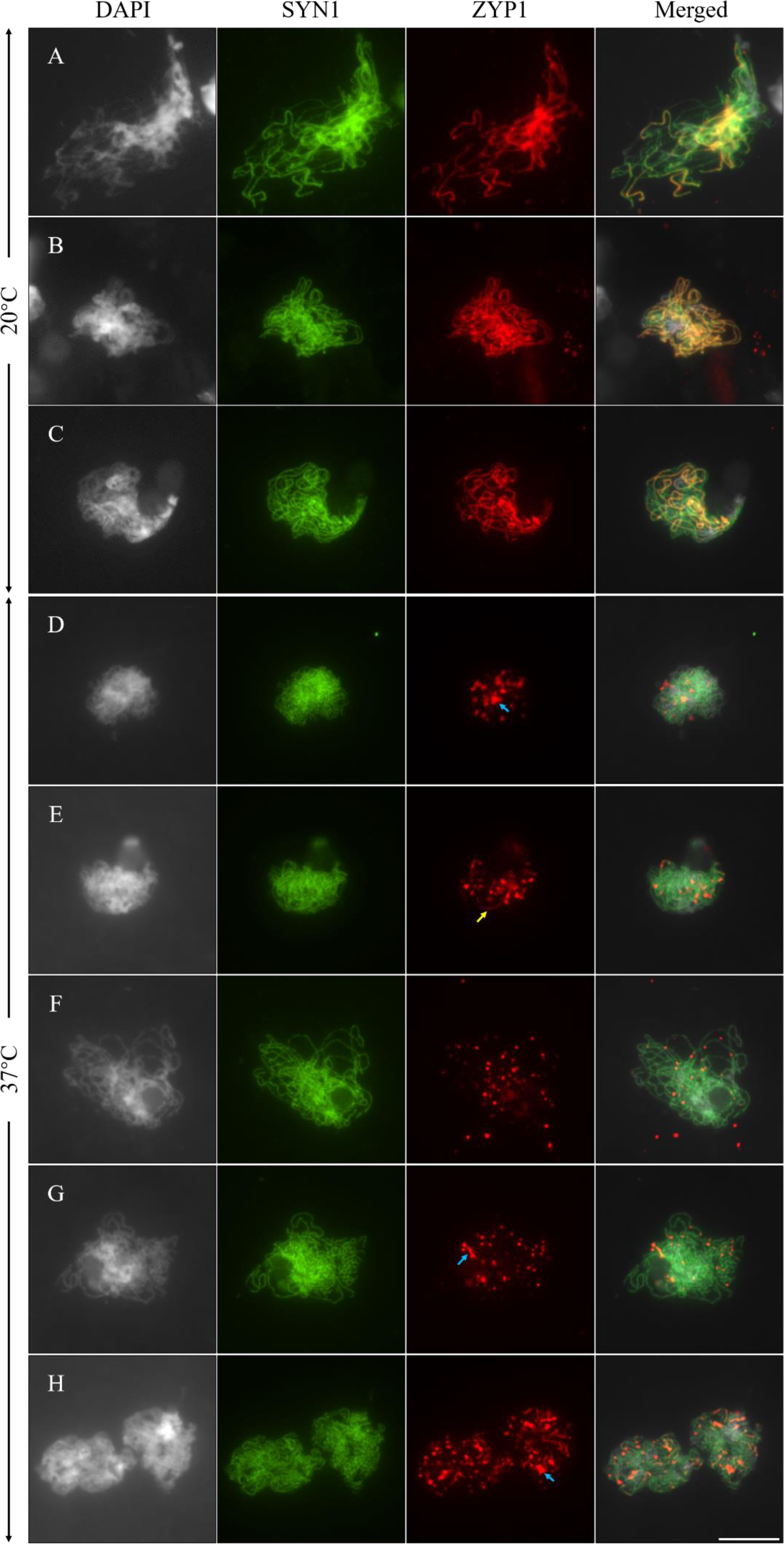
Immunolocalization of SYN1 and ZYP1 in meiocytes of autotetraploid Col-0 plants. A-C, Zygotene- (A), middle pachytene- (B) and late pachytene-staged (C) meiocytes in control plants. D-H, Early zygotene- (D), middle zygotene- (E), late zygotene- (F), middle pachytene- (G) and late pachytene-staged (H) meiocytes in heat-stress plants. Yellow arrow indicates fragmented ZYP1 signals. Blue arrows indicate aggregated and/or enlarged ZYP1 foci. Scale bar = 10 μm.

## Discussion

WGD is a conserved phenomenon that contributes to genomic diversity and speciation in higher plants (Comai, 2005; Dubcovsky and Dvorak, 2007; Ren et al., 2018; te Beest et al., 2012; Van de Peer et al., 2020). Additional copies of genome, however, increase the complexity for homologous chromosomes to pair and synapse (Lloyd and Bomblies, 2016; Svačina et al., 2020; Yant et al., 2013). In our study, the autotetraploid Col-0 plants generate a low but consistently-detectable rate (∼4.53%) of aberrant meiotic products under normal temperature conditions (Fig. 1, 2 and 3), suggesting that meiotic alterations naturally occur in these autotetraploid Arabidopsis plants. The regularly formed spindle and phragmoplast microtubule arrays in the autotetraploid Col-0 plants under control temperature (Fig. 3) indicate that the meiotic defects are not caused by alterations in microtubular cytoskeleton and cytokinesis, but are probably induced by the alterations in earlier meiosis processes, e.g. improper chromosome behaviors with a resultant impacted CO formation. Chromosome spreading analysis confirmed this which showed incomplete and/or irregular pairing and synapsis at pachytene stage (Fig. 4B-D). The observation of regions of co-aligned, but un-synapsed axes suggest that the synapsis defects are (or at least in part) independent of defects in chromosome pairing (Capilla-Pérez et al., 2021; France et al., 2021). At the same time, we found that the autotetraploid Arabidopsis plants primarily generate diakinesis PMCs that contain five tetravalents (Fig. 4). This phenotype supports the opinion that autotetraploid plants, in contrast to allotetraploids, preferentially undergo synapsis between four homologous chromosomes (Braz et al., 2021; Lloyd and Bomblies, 2016; Svačina et al., 2020). The unbalanced chromosome segregation at anaphase I could result from the difficulties in multivalent resolving in the meiocytes with a mixed existence of tetravalents and bivalents, as well as the ten bivalents (Yant et al., 2013). The autotetraploid Col-0 plants that we used here was colchicine treatment-induced, which is very typical of newly formed (neo)-autotetraploids that is believed to be less stable than the evolution-derived autotetraploids; e.g. *A. lyrate* and *A. arenosa* (Henry et al., 2014; Lloyd and Bomblies, 2016; Yant et al., 2013). Therefore, the meiotic defects we observed are probably the effects of polyploidization without natural selection.

Under heat stress, as other plant species, the autotetraploid Col-0 plants exhibit a severe organization of phragmoplast and RMAs at the end of meiosis (Fig. 3I, K and L), indicating that microtubular cytoskeleton is a prominent targeted by high temperatures (De Storme and Geelen, 2020; Lei et al., 2020; Mai et al., 2019; Wang et al., 2017). Nevertheless, heat-induced polyad formation, and the disrupted RMA configuration could be a secondary effort from the impaired spindle and/or a univalent-induced mis-segregation of chromatids at MII, which normally occur in mutants with a defective MR (Bai et al., 1999; Shi et al., 2021; Xue et al., 2019). Besides, in heat-stressed autotetraploid Arabidopsis, a high frequency of univalents is induced and the abundance of HEI10 is lowered, indicating a largely suppressed CO formation. Subsequent cytological analysis on key MR events suggested that the high temperatures interfere with MR in the autotetraploid Arabidopsis plants probably by impacting DSB formation, and by destabilizing chromosome axis and SC. Genome duplication has been suggested to provide increased tolerance to environmental stresses (Folk et al., 2020; Lourkisti et al., 2020; Rao et al., 2020). However, in contrast to the diploid Arabidopsis Col-0 plants, we here showed that the neo-autotetraploid Col-0 generates a higher frequency of aberrant meiotic products under high temperatures (Fig. 1; ∼92.5% at 37°C) (Lei et al., 2020). Meanwhile, we recorded a higher rate of destabilized loading of ASY1 and ASY4 in the heat-stressed autotetraploid Col-0 plants (80.45% defective ASY1 and 80.15% defective ASY4 in autotetraploid Col-0) (Ning et al., 2021), which represents enhanced sensitivity of axis and SC formation to increased temperature. These facts suggest that meiosis programs, especially for axis organization and chromosome dynamics are more unstable under high temperatures in neo-tetraploid *Arabidopsis thaliana*.

The significantly reduced abundance of γH2A.X and DMC1 in autotetraploid Col-0 plants under high temperatures indicated an interfered DSB generation (Fig. 6). This phenomenon is similar with what has been reported in the diploid Arabidopsis (Ning et al., 2021), which suggests that a duplicated genome does not change the threshold of DSB formation system in *Arabidopsis thaliana*. In *Saccharomyces cerevisiae*, DSB repair defect is more pronounced under lower temperature (Pohl and Nickoloff, 2008). The negative impact of high temperatures on DSB formation thus is likely conserved among eukaryotes. The attempt to reveal how heat stress influences DSB formation has been tried by examining the expression of key DSB generation-involved factors in the diploid Arabidopsis, which showed that the transcripts of neither *SPO11-1* nor *PRD1*, *2* and *3* are sensitive to the increased temperature. Therefore, heat stress induces DSB reduction not likely by impacting the DSB formation machineries at the mRNA levels. In multiple species, the activity of phosphatidylinositol 3 kinase-like (PI3K) protein kinase Ataxia-Telangiectasia Mutated (ATM), which undertakes a conserved function in sensing DNA damage and subsequently evoking DSB repair events (reviewed by (Paull, 2015)), is negatively correlated with DSB abundance (Carballo et al., 2013; Garcia et al., 2015; Joyce et al., 2011; Lange et al., 2011; Li and Yanowitz, 2019; Mohibullah and Keeney, 2017; Zhang et al., 2011). A recent report in Arabidopsis suggested that ATM limits DSB formation by restricting the accumulation of SPO11-1 on chromatin, with the *atm* mutant having increased amount of SPO11-1 (Kurzbauer et al., 2021). Supportively, the *atm* mutation increases the foci of recombinases RAD51 and DMC1; meanwhile, it enhances the chromosome fragmentation in the *atcom1* mutant, which is defective for DSB processing (Kurzbauer et al., 2021; Uanschou et al., 2007). We have previously identified an elevated expression of *ATM* in heat-stressed diploid Arabidopsis plants (Ning et al., 2021). Taken together, it is possible that high temperatures interfere with DSB formation with associated induction of univalents by activating *ATM*. To this end, a decreased SPO11-1 abundance should occur in heat-stressed wild-type plants, and the *atm* mutant may exhibit higher DSB threshold to increased temperatures. Further examination on SPO11 dynamics under increased temperature thus is of necessity to verify the hypothesis.

In both diploid and autotetraploid Arabidopsis, we observed an interfered configuration of ASY1 and ASY4 on chromosomes (Fig. 7 and 9) (Ning et al., 2021). At the same time, fragmented and/or disrupted ZYP1 loading, and multilayer-SC structures occurs in both the diploid and autotetraploid Arabidopsis under high temperatures (Fig. 10) (Loidl, 1989; Morgan et al., 2017; Ning et al., 2021). These facts reveal that chromosome axis and SC are the prominent targets by high temperatures that interfere with MR. In the *syn1* mutant, the accumulation of ASY3 and ASY4 is compromised, which, however, is not the case conversely (Fig. 8) (Chambon et al., 2018; Ferdous et al., 2012; Lambing et al., 2020b). Considering that a linear configuration of ASY3 depends on a functional ASY4 (Chambon et al., 2018), it is possible that SYN1, ASY4 and ASY3 act in a stepwise manner in mediating axis formation. However, since it has been evidenced by BiFC and Y2H assays that a direct interaction exists between ASY4 and ASY3 (Chambon et al., 2018), we cannot exclude the possibility that the normal loading of ASY4 also relies on the existence of ASY3 on the chromatin. Furthermore, it has been shown that ASY1 cannot be regularly loaded onto chromosomes in plants depleted with any of the axis-associated factors, which, however, in other way round are not impacted in the *asy1* mutant (Chambon et al., 2018; Chelysheva et al., 2005; Ferdous et al., 2012; Lambing et al., 2020a; Lambing et al., 2020b). This supports the notion that the assembly of SC occurs downstream of axis formation. In line with this, we observed a delayed unloading of ASY4 than ASY1 at later prophase I chromosomes (Supplement Fig. S8). Interestingly, heat-destabilized ASY1 and ASY4 signals occur at a very close frequency, which additionally co-localize on the chromosomes in both diploid and tetraploid Arabidopsis (Fig. 9) (Ning et al., 2021). The instability of lateral element of SC under high temperatures therefore is probably owing to an impacted axis formation. Since SYN1 keeps stable under the high temperatures, it is plausible that heat stress specifically targets the ASY4 and/or ASY3-mediated bridge structure that organizes and anchors the lateral element of SC to chromosome axis. Whether the aggregated and/or dotted ASY1 and ASY4 loading are somehow preferentially distributed at specific chromosome regions remain further investigation.

## Supplemental materials

The following files are available in the online version of this article.

Supplement Figure S1. FISH analysis of somatic cells in autotetraploid Col-0 plants.

Supplement Figure S2. Orcein staining of meiosis-staged flower buds in autotetraploid Col-0 plants stressed by 32°C.

Supplement Figure S3. Meiotic cell wall formation in heat-stressed autotetraploid Col-0 plants.

Supplement Figure S4. Meiotic spread of autotetraploid Col-0 plants.

Supplement Figure S5. Quantification of meiocytes with immunostaining of ASY1 and ASY4 in autotetraploid Col-0 plants.

Supplement Figure S6. Immunolocalization of ASY1 in meiocytes of autotetraploid Col-0 plants.

Supplement Figure S7. Immunolocalization of ASY4 in meiocytes of autotetraploid Col-0 plants.

Supplement Figure S8. Co-immunolocalization of ASY1 and ASY4 in meiocytes of autotetraploid Col-0 plants.

## Materials and methods

### Plant materials and growth conditions

Autotetraploid and diploid *Arabidopsis thaliana* Columbia-0 (Col-0) plants, and the *syn1-1* (SALK_137095), *rad51* (SAIL_873_C08), *spo11-1-1* (Grelon et al., 2001) and *dmc1* (SALK_056177) (Sanchez-Moran et al., 2007) mutants were used in the study. The autotetraploid Col-0 plants were generated as reported (De Storme and Geelen, 2011). Seeds were germinated in soil for 6-8 days and seedlings were transferred to soil and cultivated with a 16 h day/8 h night, 20°C, and 50% humidity condition. For temperature treatments, young flowering plants were transferred to a humid chamber with a 16 h day/8 h night and incubated at 32 and/or 37°C, respectively, for 24 h. All the treatment started from 8:00-10:00 AM. Meiosis-staged flower buds were fixed by carnoy’s fixative or paraformaldehyde upon the finish of treatments.

### Generation of antibodies

The anti-AtSYN1 antibodies were raised in rabbits by referring to (Bai et al., 1999); the anti-AtASY1 antibodies were generated in rabbits and mouses, respectively, against the amino acid sequence SKAGNTPISNKAQPAASRES of AtASY1 conjugated to KLH; the anti-AtZYP1 antibody (rat) was generated against the amino acid sequence GSKRSEHIRVRSDNDNVQD of AtZYP1 conjugated to KLH.

### Immunolocalization of MR proteins and α-tubulin

Immunostaining of α-tubulin and MR proteins was performed as reported (Chelysheva et al., 2010; Liu et al., 2017; Wang et al., 2014). Antibodies against ZYP1 (rabbit and/or rat) (Ning et al., 2020), DMC1 (rabbit) (Ning et al., 2020) and γH2A.X (rabbit) (Lambing et al., 2020b) were diluted by 1:100; antibodies against α-tubulin (rat) (Lei et al., 2020), ASY1 (rabbit and/or mouse), ASY4 (rabbit) (Ning et al., 2020) and SYN1 (mouse) were diluted by 1:200; antibody against CENH3 (rabbit) (Abcam, 72001) was diluted by 1:400; antibody against SYN1 (rabbit) was diluted by 1:500. The secondary antibodies; i.e. Goat anti-Rabbit IgG (H+L) Cross-Adsorbed Secondary Antibody Alexa Fluor 555 (Invitrogen, A32732), Goat anti-Rabbit IgG (H+L) Highly Cross-Adsorbed Secondary Antibody Alexa Fluor Plus 488 (Invitrogen, A32731), Goat anti-Rat IgG (H+L) Cross-Adsorbed Secondary Antibody, Alexa Fluor 555 (Invitrogen, A21434), Goat anti-Rat IgG (H+L) Cross-Adsorbed Secondary Antibody, Alexa Fluor 488 (Invitrogen, A11006) and Goat anti-Mouse IgG (H+L) Highly Cross-Adsorbed Secondary Antibody, Alexa Fluor Plus 488 (Invitrogen, A32723) were diluted to 10 µg/mL.

### Cytology and fluorescence in situ hybridization

Meiotic chromosome behaviors were analyzed by performing chromosome spreading using meiosis-staged flower buds fixed at least 24 h by carnoy’s fixative. Flower buds were washed twice by distilled water and once in citrate buffer (10 mM, pH = 4.5), and were incubated in digestion enzyme mixture (0.3% pectolyase, 0.3% cellulase and 0.3% cytohelicase) in citrate buffer (10 mM, pH = 4.5) at 37°C in a moisture chamber for 2.5-3.5 h. Subsequently, 6-8 digested buds were washed in distilled water, and were transferred to a glass slide squashed in a small amount (4-5 μL) of distilled water followed by adding two rounds of 10 μL precooled 60% acetic acid. The samples were stirred gently on a hotplate at 45°C for 1-2 min, which thereafter was flooded with precooled carnoy’s fixative. The slides were subsequently air dried for 10 min, and were stained by adding 8 μL DAPI (10 μg/mL) in Vectashield antifade mounting medium, mounted with a coverslip, and sealed by nail polish. Tetrad analysis by orcein and/or aniline blue staining was performed by referring to (Lei et al., 2020; Ning et al., 2021). FISH assay was performed by referring to (Lei et al., 2020).

### Microscopy and quantification of fluorescent foci

Bright-field images and DAPI-stained meiotic chromosomes were pictured using a M-Shot ML31 microscope equipped with a MS60 camera. Aniline blue staining of meiotic cell walls, and immunolocalization of α-tubulin and MR-related proteins were analyzed on an Olympus IX83 inverted fluorescence microscope equipped with a X-Cite lamp and a Prime BSI camera. Image processing and quantification of fluorescent foci were conducted as previously reported (Ning et al., 2021).

## Author contribution

H.Q.F. performed most of the experiments; K.Y., X.H.Z. and J.Y.Z. performed meiotic spread analysis; E.I.E. performed FISH experiment; H.L., J.X., C.L.C. and G.H.Y. contributed to data analysis; C.W. analyzed ɤH2A.X and DMC1 foci; B.L. conceived the project, analyzed data, and wrote the manuscript.

## Acknowledgement

The authors thank Dr. Jing Li (Huazhong Agricultural University) and Dr. Yingxiang Wang (Fudan University) for kindly providing the autotetraploid Col-0 seeds, and the anti-HEI10 (rabbit) antibody, respectively. They appreciate Dr. Wojtek Pawlowski (Cornell University) for the discussion and suggestions while collecting data. Especially, the authors appreciate Dr. Andrew Lloyd (IBERS) for critical review and comments on the manuscript prior to submission.

